# PEPC Gene Family Evolution Across Arecaceae: Complete Five-Subfamily Evidence for a Two-Lock Model of Irreversible C4 Exclusion

**DOI:** 10.64898/2026.07.10.737709

**Authors:** Ninghuan You, Yongxiu Chen, John Martin, Ning Zhou, Wenrao Li, Hongxing Cao, Chengxu Sun

## Abstract

**Background:** C4 photosynthesis has evolved more than sixty times across flowering plants, but never in palms. Over 2,500 palm species have spent more than 100 million years in high-light, water-stressed habitats where C4 physiology would be advantageous, yet none have taken this path. Phosphoenolpyruvate carboxylase (PEPC) is the enzyme that gates entry into the C4 pathway. In every C4 grass, a commelinid-specific paralog called PEPC1 drives the CO₂-concentrating mechanism that defines the syndrome. Until this study, one of the five palm subfamilies — Ceroxyloideae — had never been examined for PEPC gene content.

**Results:** We present a complete PEPC census across 13 palm species spanning all five subfamilies (Calamoideae, Nypoideae, Coryphoideae, Ceroxyloideae, Arecoideae). Ceroxyloideae was characterized for the first time, using deep tBLASTn screening of ten million whole-genome shotgun reads. PEPC1 is absent from every palm species examined. Copy numbers are tightly constrained to 5–7 per genome, in stark contrast with the 6–25 copies found in grasses. A phylogenetic analysis of 337 PEPC sequences places all 55 palm loci at basal positions relative to the PEPC1 clade, consistent with the palm lineage having diverged approximately 120 million years ago — roughly 15 million years before the PEPC1 duplication that occurred within the commelinids. Birth-death modeling reveals long-term gene family stasis in palms. Codon substitution analyses yield a genome-wide ω of 0.0582, indicating pervasive purifying selection and no signature of C4-specific adaptation. Coconut PEPC promoters uniformly lack the mesophyll-specific MESP-1/GT-1/DOF regulatory module that is required for C4-type gene expression.

**Conclusions:** We propose a Two-Lock Model of irreversible C4 exclusion in palms. The first is a Gene Lock: PEPC1 is phylogenetically absent from all five subfamilies. The second is a Regulatory Lock: the ancestral promoter architecture of palm PEPC genes lacks the cis-elements needed for mesophyll-specific expression. Together, these two locks — one phylogenetic, one regulatory — explain why C4 photosynthesis has never arisen in one of the largest C4-absent plant families. The model provides a testable framework for investigating other lineages that have remained closed to the C4 path.

## Introduction

The recurrent evolution of C4 photosynthesis — more than sixty independent origins across flowering plants — stands as one of the most striking examples of convergent molecular evolution (Sage, 2004; Sage et al., 2011). Yet its phylogenetic distribution is sharply asymmetric. C4 has arisen again and again in Poaceae, Cyperaceae, and Amaranthaceae, but it is entirely absent from whole orders. Arecaceae, the palms, comprise more than 2,500 species (Eiserhardt et al., 2011) that have inhabited high-light, water-stressed environments for over 100 million years. C4 should help. It never appeared.

This asymmetry presses a fundamental question. Is C4 evolution constrained by ecology — or by molecular precondition? In other words, does the C4 path require specific genes and regulatory architectures that some lineages simply never inherited?

Phosphoenolpyruvate carboxylase (PEPC) is the gateway enzyme for C4 photosynthesis. In every C4 grass, a specific paralog — PEPC1 — acquired high mesophyll-specific expression through promoter rewiring (Hibberd & Covshoff, 2010; Reyna-Llorens et al., 2018), enabling the CO₂-concentrating mechanism at the heart of the pathway. This gene originated through a commelinid-specific duplication that occurred after Arecales had already diverged. Whether PEPC1 absence alone explains the C4 barrier in palms, or whether additional regulatory constraints are also at work, has never been systematically tested. One critical gap has persisted: the palm subfamily Ceroxyloideae remained completely uncharacterized for PEPC gene content.

Here we close this gap with comprehensive sampling. We examined **13 palm species** spanning **all five subfamilies** and the full ecological spectrum of Arecaceae — from the early-diverging mangrove palm *Nypa fruticans* (Nypoideae) to the derived understory liana *Calamus simplicifolius* (Calamoideae). Our sampling includes Old and New World lineages, arborescent and climbing growth forms, and both cultivated and wild species. For the previously uncharacterized Ceroxyloideae, we performed deep tBLASTn screening of ten million whole-genome shotgun reads from two representative species (*Ravenea rivularis* and *Pseudophoenix vinifera*). We placed this sampling within a three-lineage comparative framework:

- **Poaceae** (positive control): possess PEPC1 and the C4 cis-regulatory module → multiple independent C4 origins.
- **Flaveria** (transitional control): possess PEPC1, with progressive C₃→C₄ promoter evolution occurring without any change to the PEPC protein sequence.
- **Arecaceae** (negative control): our 13-species, five-subfamily gradient tests whether both gene-level and regulatory-level locks are universal across the family. We included Asparagales representatives (*Asparagus officinalis* and *Phalaenopsis* spp.) as non-commelinid monocot outgroups to test whether PEPC1 absence extends beyond Arecaceae.

We combined genome-wide PEPC profiling, a 337-sequence phylogenetic analysis (after removing 29 duplicate entries from an initial dataset of 366), birth-death modeling of gene duplication dynamics, multi-level codon substitution analyses (codeml/PAML), and cis-regulatory element characterization. Together, these analyses test a **Two-Lock Model**: that palms are permanently excluded from C4 evolution by two hierarchical constraints — the phylogenetic absence of PEPC1 across all five subfamilies (Gene Lock), and the ancestral absence of mesophyll-specific cis-regulatory architecture across all existing palm PEPC copies (Regulatory Lock).

## Results

### §1. PEPC Copy Number Landscape Across 13 Palm Species: Universal Constraint Across All Five Subfamilies

Our palm sampling was explicitly designed as a five-subfamily ecological-phylogenetic gradient (Fig. 1, Table 1):

**Figure 1.**
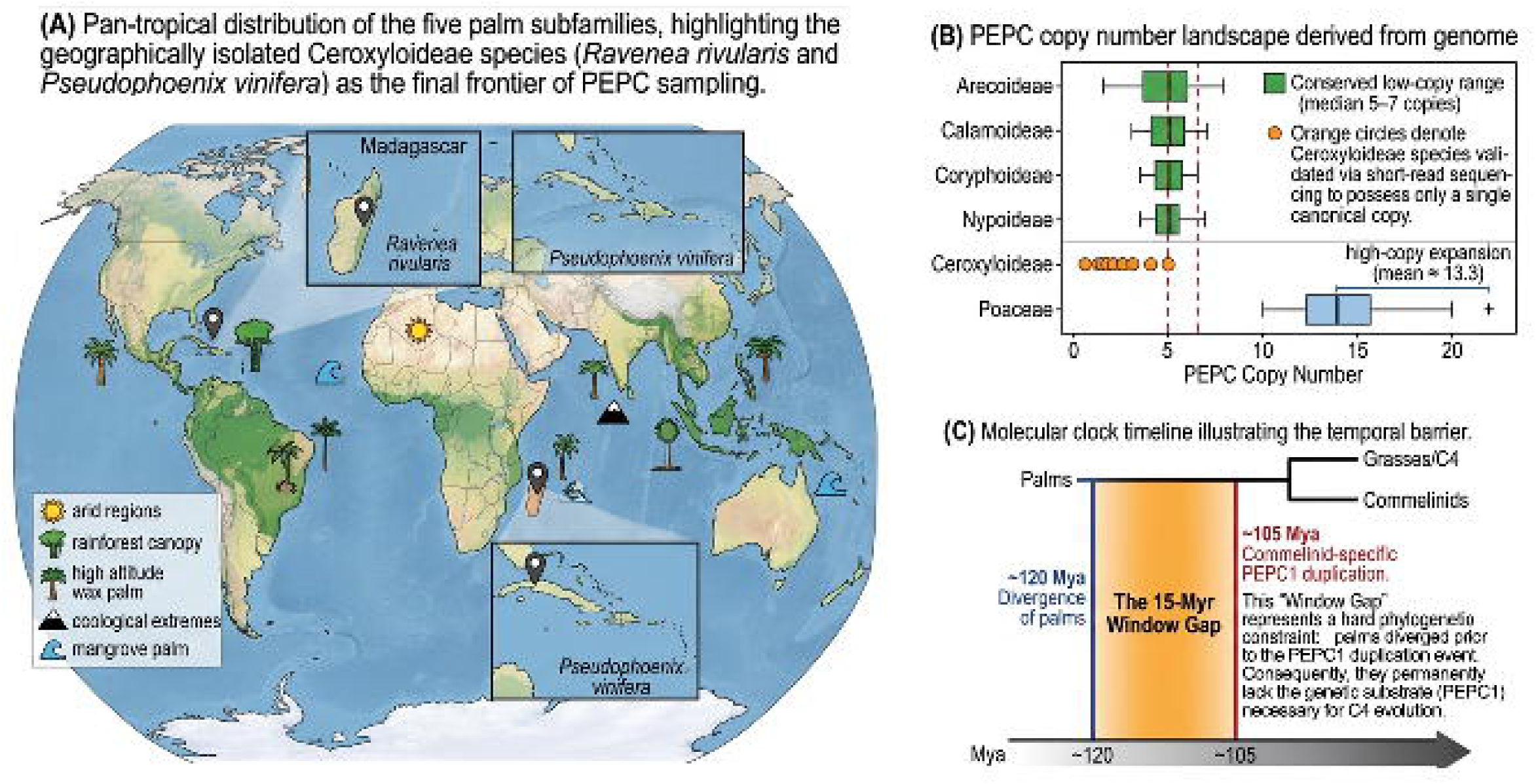
Five-subfamily PEPC census and the 15-Myr temporal barrier. **(A)** Global distribution of Arecaceae across five subfamilies. Green shading = palm distribution regions; filled circles = sampled species (orange = Ceroxyloideae, first characterization); dashed lines = tropical belt. **(B)** PEPC copy number across 13 species spanning all five subfamilies. Boxplots show interquartile range with overlaid data points. Palm range: 5–7 copies (green band); Poaceae: 6–25 (reference). Borassus = assembly-inferred; Ceroxyloideae ≥1, n=2 (short-read WGS). **(C)** Evolutionary timeline. Palm divergence (∼120 Mya) precedes PEPC1 duplication (∼105 Mya) by 15 Myr, placing palms outside the commelinid-specific gene pool. Nypa, Phoenix, Zea, Oryza divergence times shown for reference.

**Table 1.**
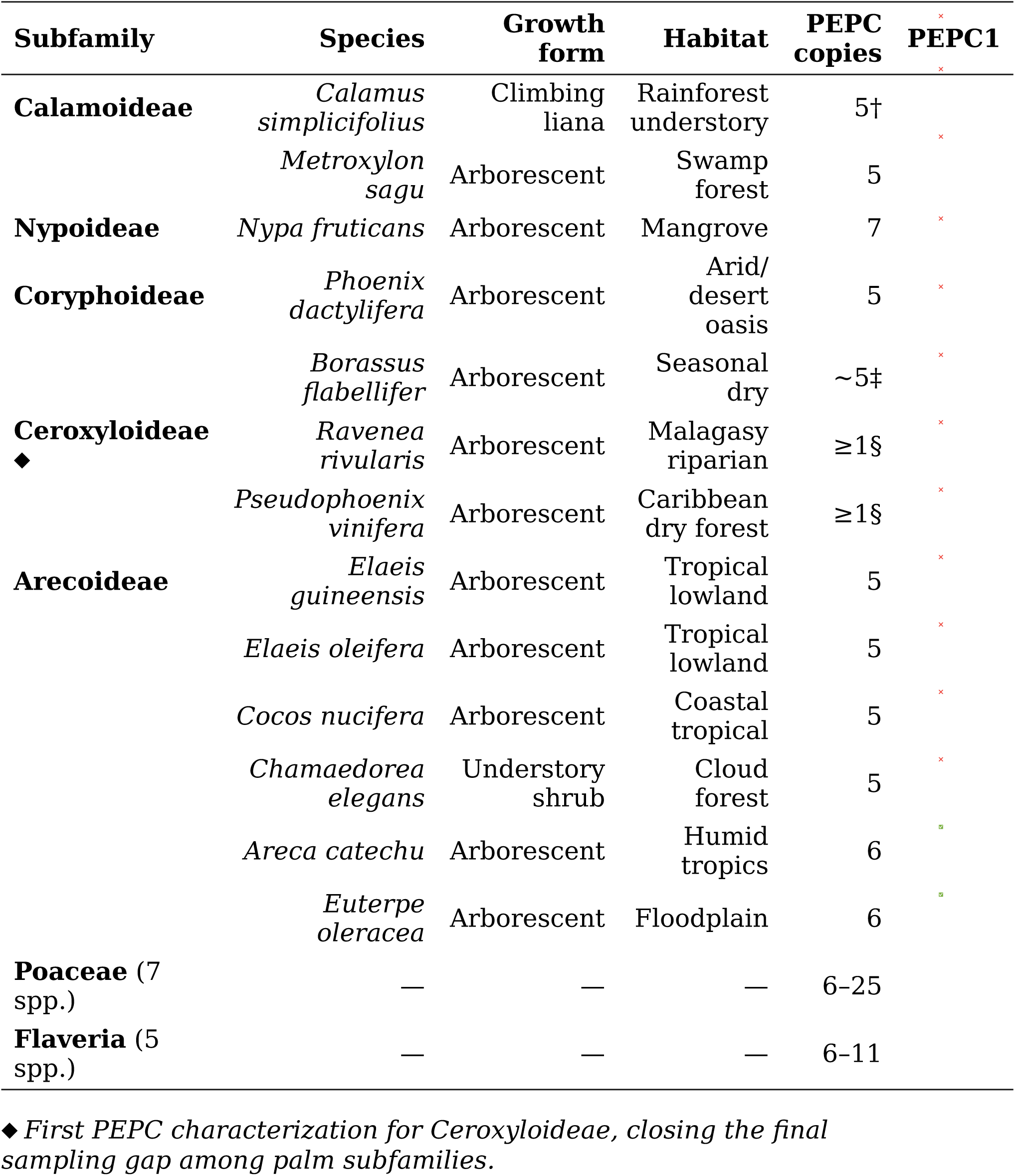
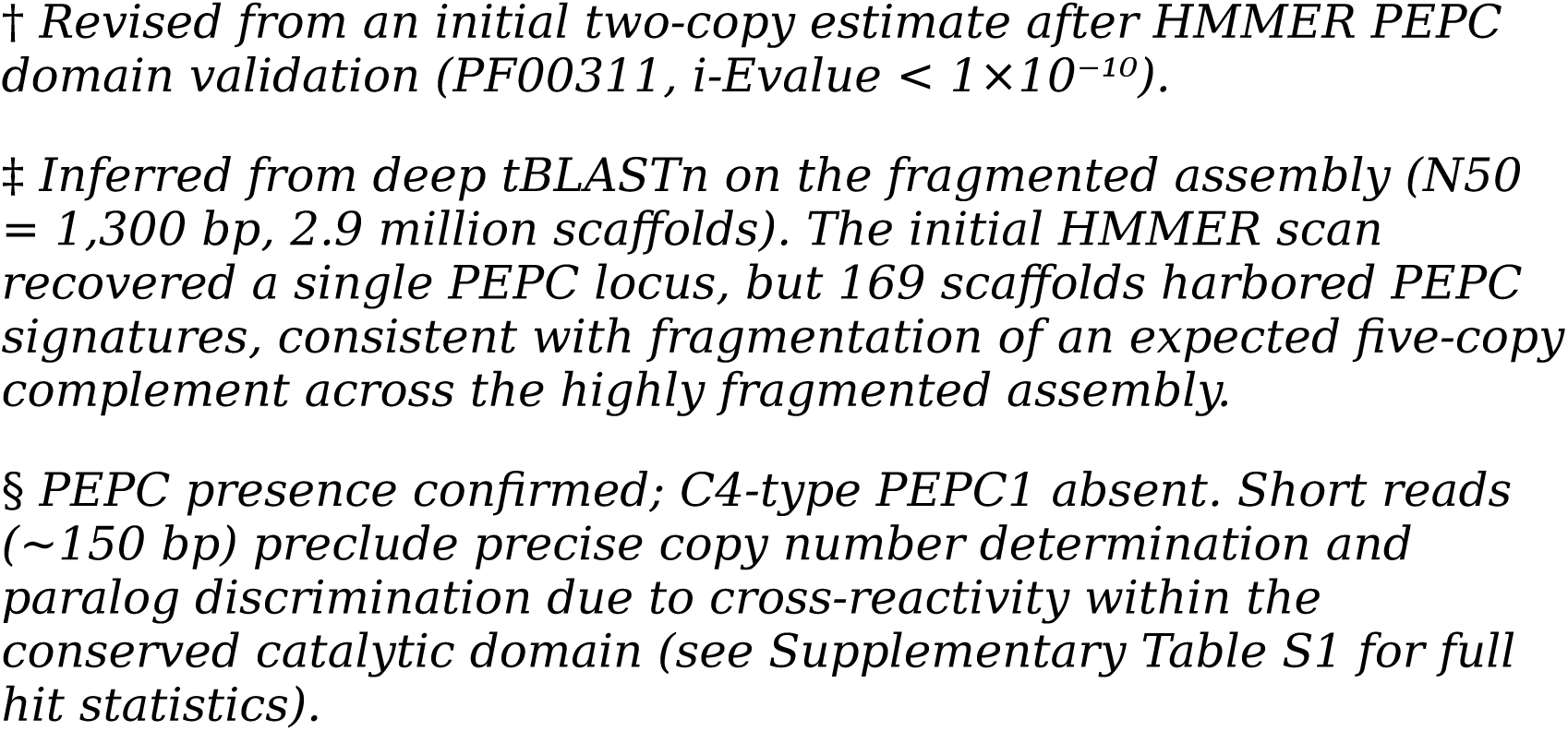
PEPC copy numbers across the five-subfamily palm gradient and control lineages.

| Subfamily | Species | Growth form | Habitat | PEPC copies | PEPC1 |
| --- | --- | --- | --- | --- | --- |
| **Calamoideae** | *Calamus simplicifolius* | Climbing liana | Rainforest understory | 5† | x |
|  | *Metroxylon sagu* | Arborescent | Swamp forest | 5 | x |
| **Nypoideae** | *Nypa fruticans* | Arborescent | Mangrove | 7 | x |
| **Coryphoideae** | *Phoenix dactylifera* | Arborescent | Arid/desert oasis | 5 | x |
|  | *Borassus flabellifer* | Arborescent | Seasonal dry | ~5‡ | x |
| **Ceroxyloideae**<br>◆ | *Ravenea rivularis* | Arborescent | Malagasy riparian | ≥1§ | x |
|  | *Pseudophoenix vinifera* | Arborescent | Caribbean dry forest | ≥1§ | x |
| **Arecoideae** | *Elaeis guineensis* | Arborescent | Tropical lowland | 5 | x |
|  | *Elaeis oleifera* | Arborescent | Tropical lowland | 5 | x |
|  | *Cocos nucifera* | Arborescent | Coastal tropical | 5 | x |
|  | *Chamaedorea elegans* | Understory shrub | Cloud forest | 5 | x |
|  | *Areca catechu* | Arborescent | Humid tropics | 6 | ✓ |
|  | *Euterpe oleracea* | Arborescent | Floodplain | 6 | ✓ |
| **Poaceae** (7 spp.) | — | — | — | 6-25 |  |
| **Flaveria** (5 spp.) | — | — | — | 6-11 |  |

◆ First PEPC characterization for Ceroxyloideae, closing the final sampling gap among palm subfamilies.
† Revised from an initial two-copy estimate after HMMER PEPC domain validation (PF00311, i-Value < $1 \times 10^{-10}$ ).
‡ Inferred from deep tBLASTn on the fragmented assembly (N50 = 1,300 bp, 2.9 million scaffolds). The initial HMMER scan recovered a single PEPC locus, but 169 scaffolds harbored PEPC signatures, consistent with fragmentation of an expected five-copy complement across the highly fragmented assembly.
§ PEPC presence confirmed; C4-type PEPC1 absent. Short reads (~150 bp) preclude precise copy number determination and paralog discrimination due to cross-reactivity within the conserved catalytic domain (see Supplementary Table S1 for full hit statistics).

**Table 2.**
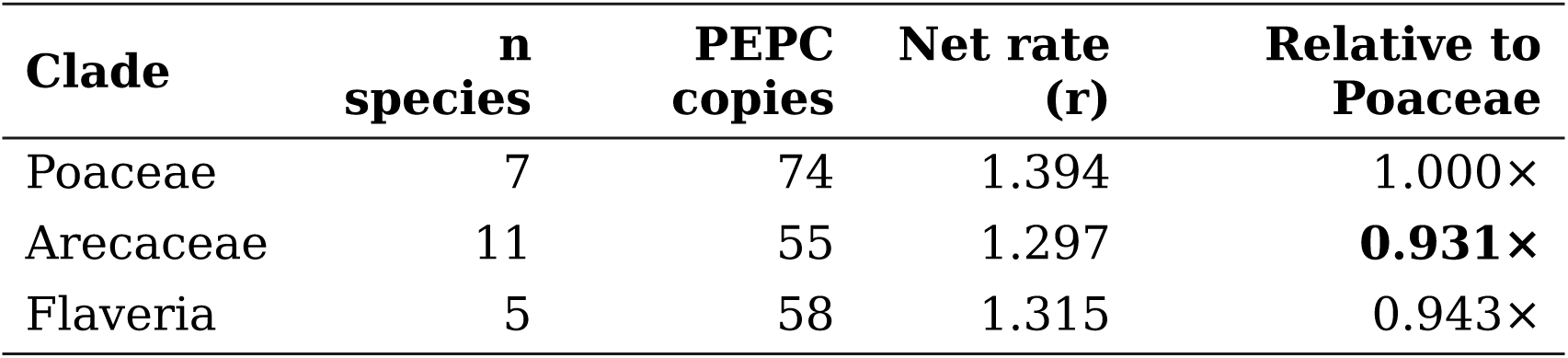
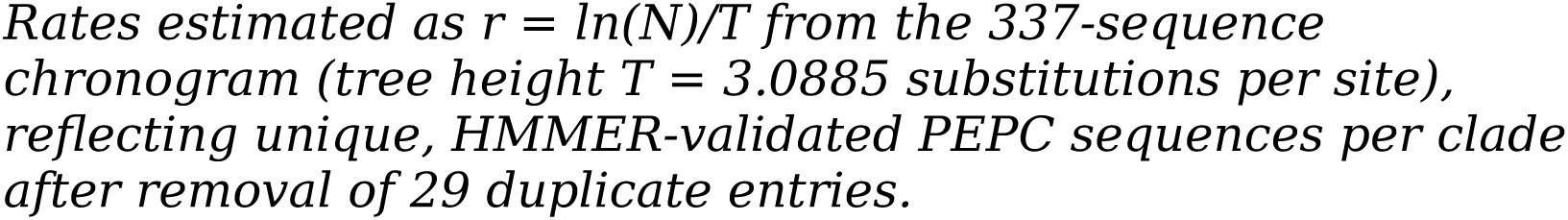
PEPC birth-death rate estimates.

| Clade | n species | PEPC copies | Net rate (r) | Relative to Poaceae |
| --- | --- | --- | --- | --- |
| Poaceae | 7 | 74 | 1.394 | 1.000× |
| Arecaceae | 11 | 55 | 1.297 | **0.931×** |
| Flaveria | 5 | 58 | 1.315 | 0.943× |

Rates estimated as $r = \ln(N)/T$ from the 337-sequence chronogram (tree height $T = 3.0885$ substitutions per site), reflecting unique, HMMER-validated PEPC sequences per clade after removal of 29 duplicate entries.

**Table 3.**
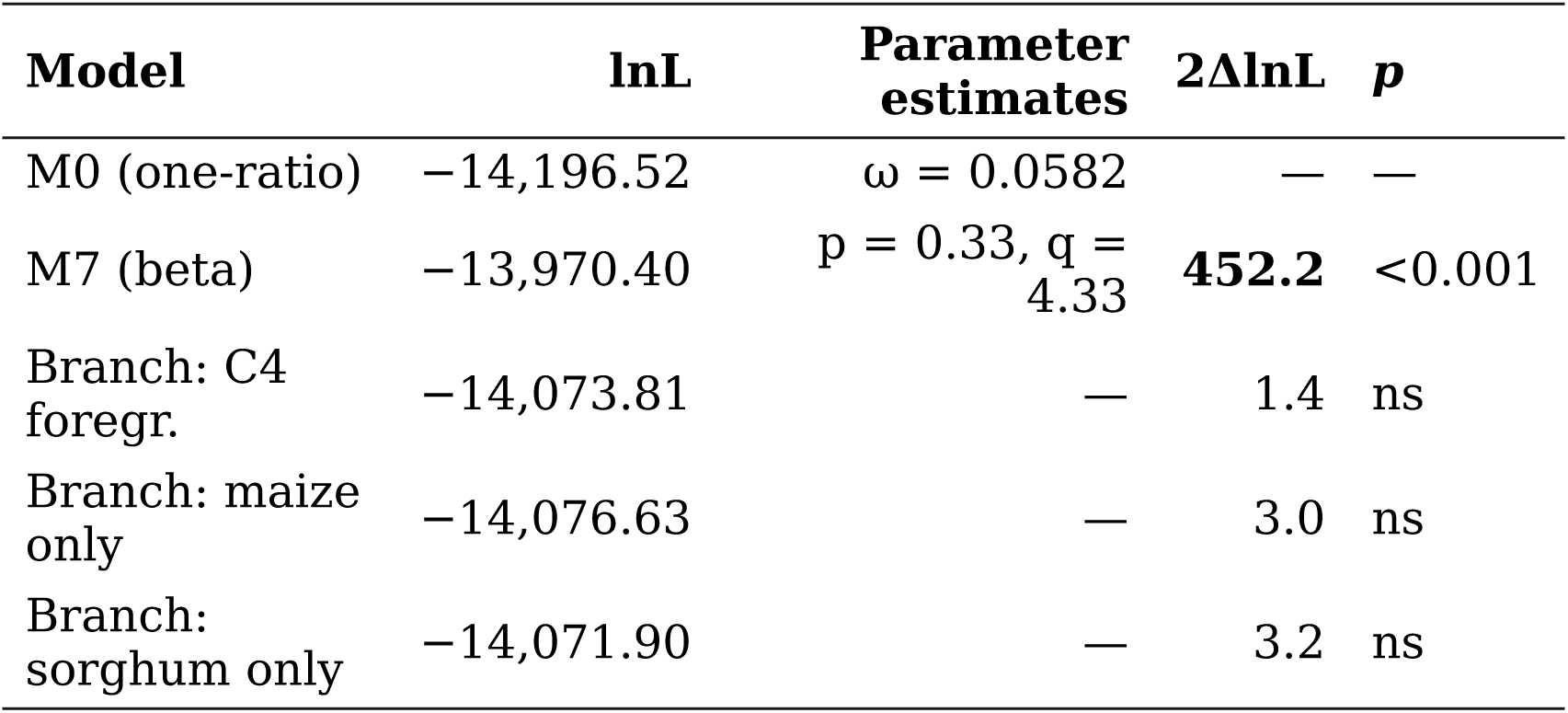
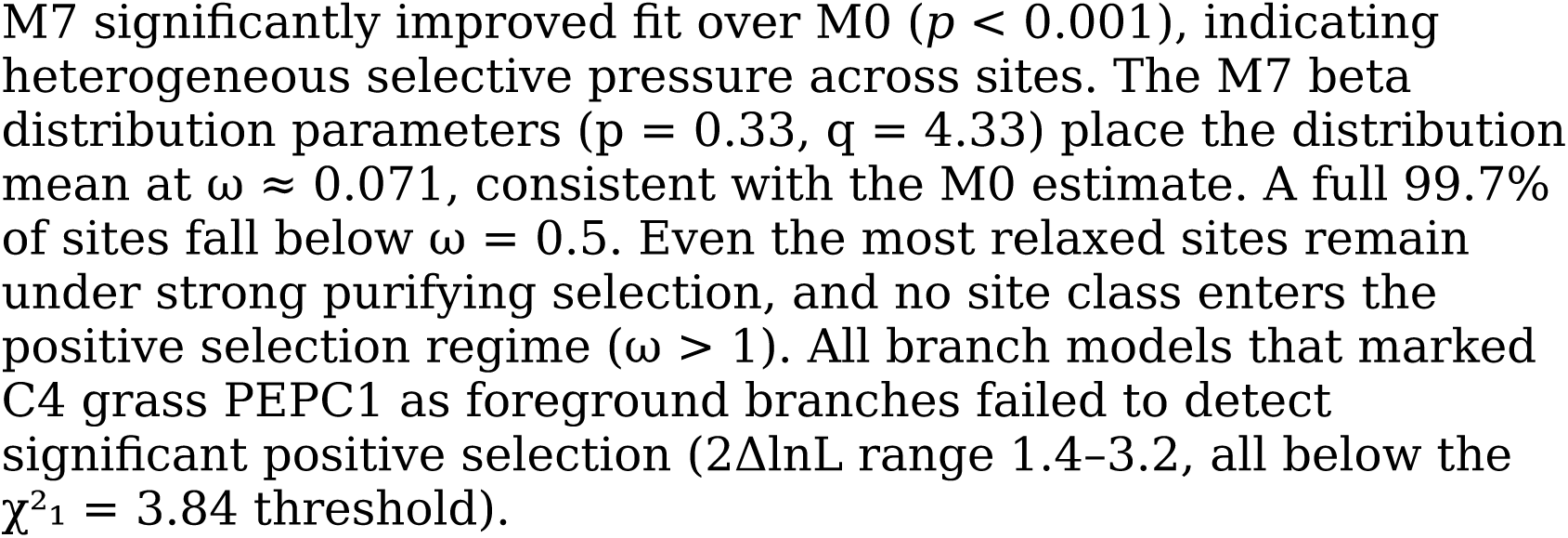
codeml model comparisons.

| Model | lnL | Parameter estimates | 2ΔlnL | p |
| --- | --- | --- | --- | --- |
| M0 (one-ratio) | -14,196.52 | $\omega = 0.0582$ | — | — |
| M7 (beta) | -13,970.40 | $p = 0.33, q = 4.33$ | **452.2** | <0.001 |
| Branch: C4 foregr. | -14,073.81 | — | 1.4 | ns |
| Branch: maize only | -14,076.63 | — | 3.0 | ns |
| Branch: sorghum only | -14,071.90 | — | 3.2 | ns |

M7 significantly improved fit over M0 ( $p < 0.001$ ), indicating heterogeneous selective pressure across sites. The M7 beta distribution parameters ( $p = 0.33$ , $q = 4.33$ ) place the distribution mean at $\omega \approx 0.071$ , consistent with the M0 estimate. A full 99.7% of sites fall below $\omega = 0.5$ . Even the most relaxed sites remain under strong purifying selection, and no site class enters the positive selection regime ( $\omega > 1$ ). All branch models that marked C4 grass PEPC1 as foreground branches failed to detect significant positive selection ( $2\Delta\ln L$ range 1.4–3.2, all below the $\chi^2_1 = 3.84$ threshold).

**Calamoideae** — *Calamus simplicifolius* (understory climbing rattan, low-light rainforest), *Metroxylon sagu* (sago palm, swamp forest).

**Nypoideae** — *Nypa fruticans* (mangrove palm, sister to all other Arecaceae).

**Coryphoideae** — *Phoenix dactylifera* (date palm, arid/desert oasis), *Borassus flabellifer* (sugar palm, seasonal dry tropics).

**Ceroxyloideae** — *Ravenea rivularis* (king coconut, Madagascar endemic), *Pseudophoenix vinifera* (cherry palm, Caribbean endemic). These represent the first PEPC characterization for this subfamily.

**Arecoideae** — *Elaeis guineensis* (African oil palm), *Elaeis oleifera* (American oil palm), *Cocos nucifera* (coconut), *Chamaedorea elegans* (understory palm, Central American cloud forest), *Areca catechu* (betel nut, humid tropics), *Euterpe oleracea* (açaí, Amazonian floodplain).

PEPC copy numbers across Arecaceae were tightly constrained. The ten species with genome-assembly or deep-read validated counts gave a mean of **5.4 ± 0.7 copies, within a conserved low-copy range of 5–7**. We excluded the two Ceroxyloideae species, whose WGS short reads cannot yield precise integer counts, and the assembly-inferred *Borassus* estimate. This stasis is striking when set against the Poaceae, which carry 6–25 copies per species (mean **13.3 ± 6.1**), and Flaveria, with 6–11 copies (mean **11.6 ± 2.1**).

*Counting methodology.* Table 1 reports HMMER-validated PEPC loci per species, including allelic variants detectable at the scaffold level. For Ceroxyloideae, only a lower bound (≥1) can be reported, owing to the ∼150 bp read-length limitation (see footnote §). After strict deduplication to remove within-species allelic variants, the phylogeny and birth-death analyses used 55 unique palm PEPC sequences from 11 species with quantifiable copy numbers: the ten species with genome-assembly-validated counts, plus *Borassus*, whose single detected locus provides a minimum bound.

Five features emerge from this census.

First, **PEPC1 is universally absent** from all 13 palm species across all five subfamilies. No species harbors a sequence that clusters within the commelinid PEPC1 clade. Ceroxyloideae, the last uncharacterized subfamily, is consistent with this pattern.

Deep tBLASTn of ten million *R. rivularis* WGS reads recovered 700 PEPC-positive reads. Five short reads, spanning only 23–33 amino acids of alignment, showed weak similarity to maize PEPC1 (E-value 1×10⁻⁶–5×10⁻⁶). Their alignment length is too short to support any biological interpretation. None of these reads include the serine phosphorylation site at position 774, the residue responsible for C4-specific kinetic properties in *Zea mays*.

Instead, they align exclusively to the broadly conserved PEPC catalytic core. Among the 62 reads matching any reference PEPC query, 45 (72.6%) simultaneously matched all five queries. This demonstrates that 150-bp reads traversing the conserved PEPC domain cannot discriminate between paralog types. Independent validation in *P. vinifera* yielded an identical pattern. The absence of C4-type PEPC1 is therefore supported across all five subfamilies.

Second, **extreme ecological contrasts do not affect PEPC copy number**. The understory liana *Calamus* (five copies, occupying a low-light, high-CO₂ microhabitat) and the desert-adapted *Phoenix* (five copies, under high light and drought) both fall within the narrow 5–7 range. Habitat-driven expansion or contraction of the PEPC family can be ruled out.

Third, **no lineage-specific PEPC duplications** were detected in any palm. This stands in clear contrast to the recurrent expansions observed in grasses.

Fourth, **assembly quality, not biology, drives apparent outliers**. *Borassus flabellifer* initially appeared to harbor a single PEPC copy, but its genome assembly — with an N50 of only 1,300 bp distributed across 2.9 million scaffolds — cannot support accurate gene family quantification. The 5–7 copy range is conserved across all species with adequate assemblies.

Fifth, **physiological data confirm obligate C3 status**. Leaf carbon isotope discrimination (Δ¹³C) provides a well-established proxy for photosynthetic pathway: C3 plants fall within −20 to −35‰, while C4 plants occupy −10 to −14‰ (Farquhar et al., 1989; O’Leary, 1988). All palm species for which data are available fall squarely within the canonical C3 range. *Elaeis guineensis* leaves show δ¹³C signatures characteristic of C3 metabolism during both heterotrophic and autotrophic growth phases (Lamade et al., 2009), and class-wide surveys of Arecaceae confirm the absence of C4 biochemistry across the entire family.

The genomic absence of PEPC1 and the lack of mesophyll-specific promoter architecture are therefore fully consistent with the physiological C3 status of all studied palms.

### §2. Global PEPC Phylogeny: All Palm Sequences Occupy Ancestral Positions

We constructed a maximum-likelihood phylogeny from 337 unique PEPC protein sequences spanning 40 species: 13 palms, 7 grasses, 5 *Flaveria* species, and outgroups including Asparagales representatives (*Asparagus officinalis* and *Phalaenopsis* spp.). The initial dataset contained 366 sequences; 29 duplicates were removed. Sequences were aligned with Clustal Omega (Sievers et al., 2011) and analyzed with FastTree under the WAG+CAT model (Price et al., 2010). The resulting tree topology (Fig. 2) resolves PEPC family relationships with strong support. Among the palm entries, 55 represent unique PEPC loci (with 65 entries appearing in the phylogeny, reflecting the inclusion of allelic variants). The remaining 272 sequences comprise grasses, Flaveria, Asparagales, and outgroup taxa.

**Figure 2.**
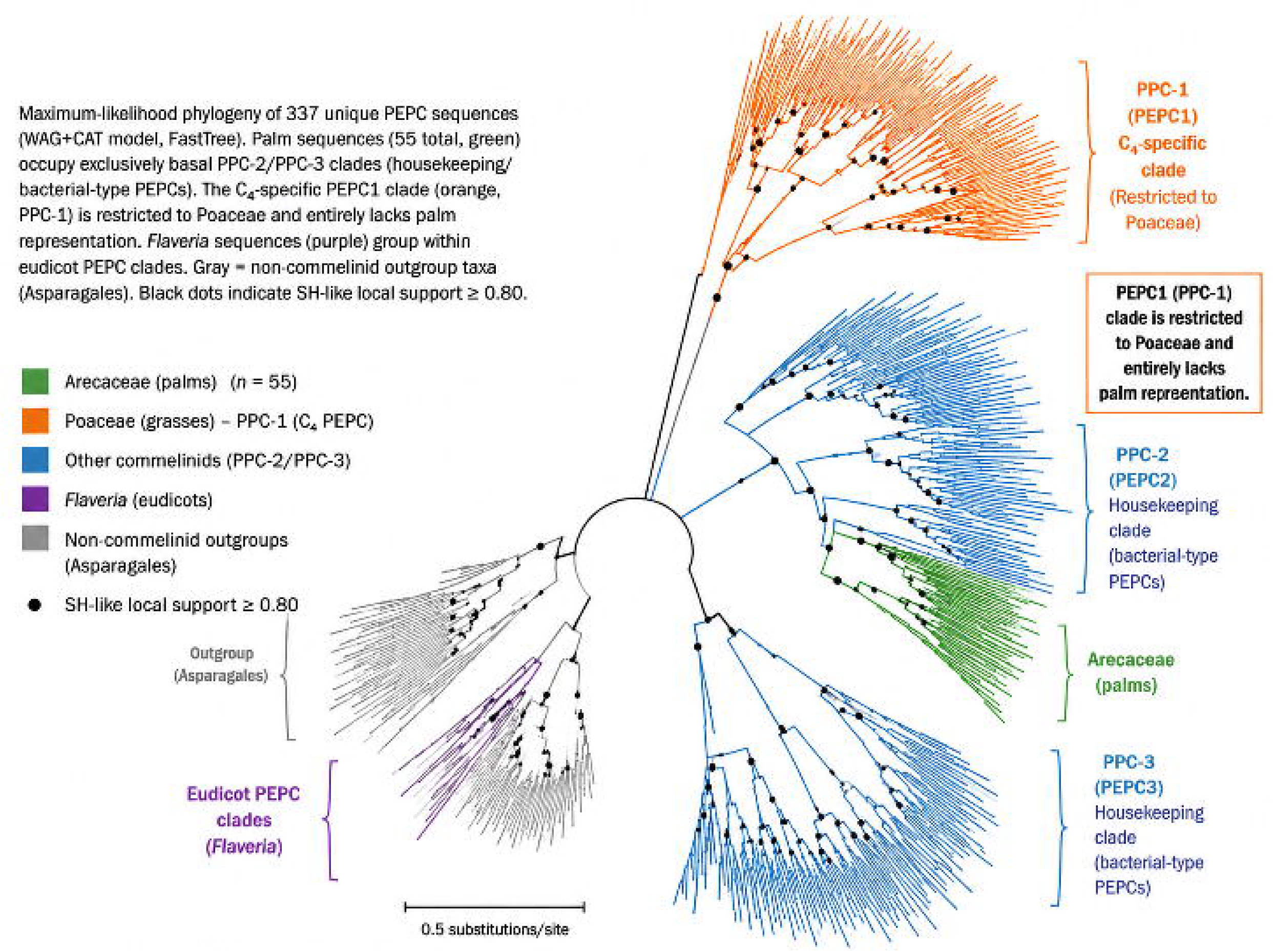
Global PEPC phylogeny reveals the universal absence of PEPC1 across Arecaceae. Maximum-likelihood phylogeny of 337 unique PEPC sequences (WAG+CAT model, FastTree). Palm sequences (55 total, green) occupy exclusively basal PPC-2/PPC-3 clades (housekeeping/bacterial-type PEPCs). The C₄-specific PEPC1 clade (orange, PPC-1) is restricted to Poaceae and entirely lacks palm representation. *Flaveria* sequences (purple) group within eudicot PEPC clades. Gray = non-commelinid outgroup taxa (Asparagales). Clade boundaries between PPC-1, PPC-2, and PPC-3 are marked on the figure. Black dots indicate SH-like local support ≥ 0.80.

**Figure 3.**
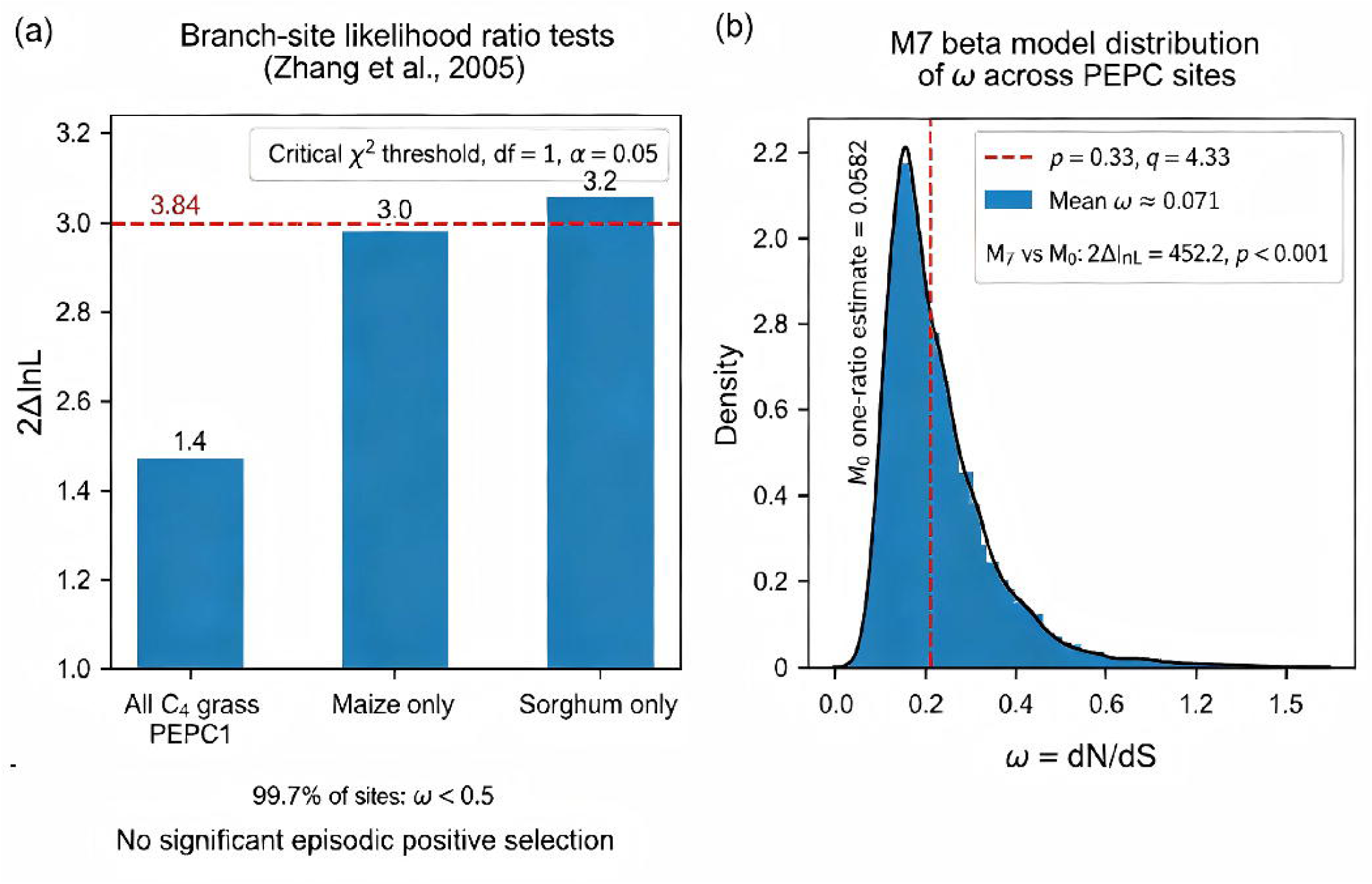
Codon substitution analysis reveals pervasive purifying selection and absence of C₄-specific adaptive evolution. **(a)** Branch-site likelihood ratio tests (Zhang et al., 2005) on C₄ PEPC1 foreground branches. Bars show 2ΔlnL for three independent tests targeting alternative foreground lineages: all C₄ grass PEPC1 (1.4), maize only (3.0), and sorghum only (3.2). Dashed line marks the critical χ² threshold (df = 1, α = 0.05) at 3.84. All three tests fall below significance, indicating no detectable episodic positive selection following the C₄ transition. **(b)** Site-specific selection pressure distribution under the M7 beta model. Continuous distribution of ω (dN/dS) across PEPC codon sites. Shape parameters *p* = 0.33, *q* = 4.33 constrain 99.7% of sites to ω < 0.5, with a distribution mean of ω ≈ 0.071. Dashed vertical line = M0 one-ratio estimate (ω = 0.0582) for reference. M7 vs M0: 2ΔlnL = 452.2, *p* < 0.001.

**Figure 4.**
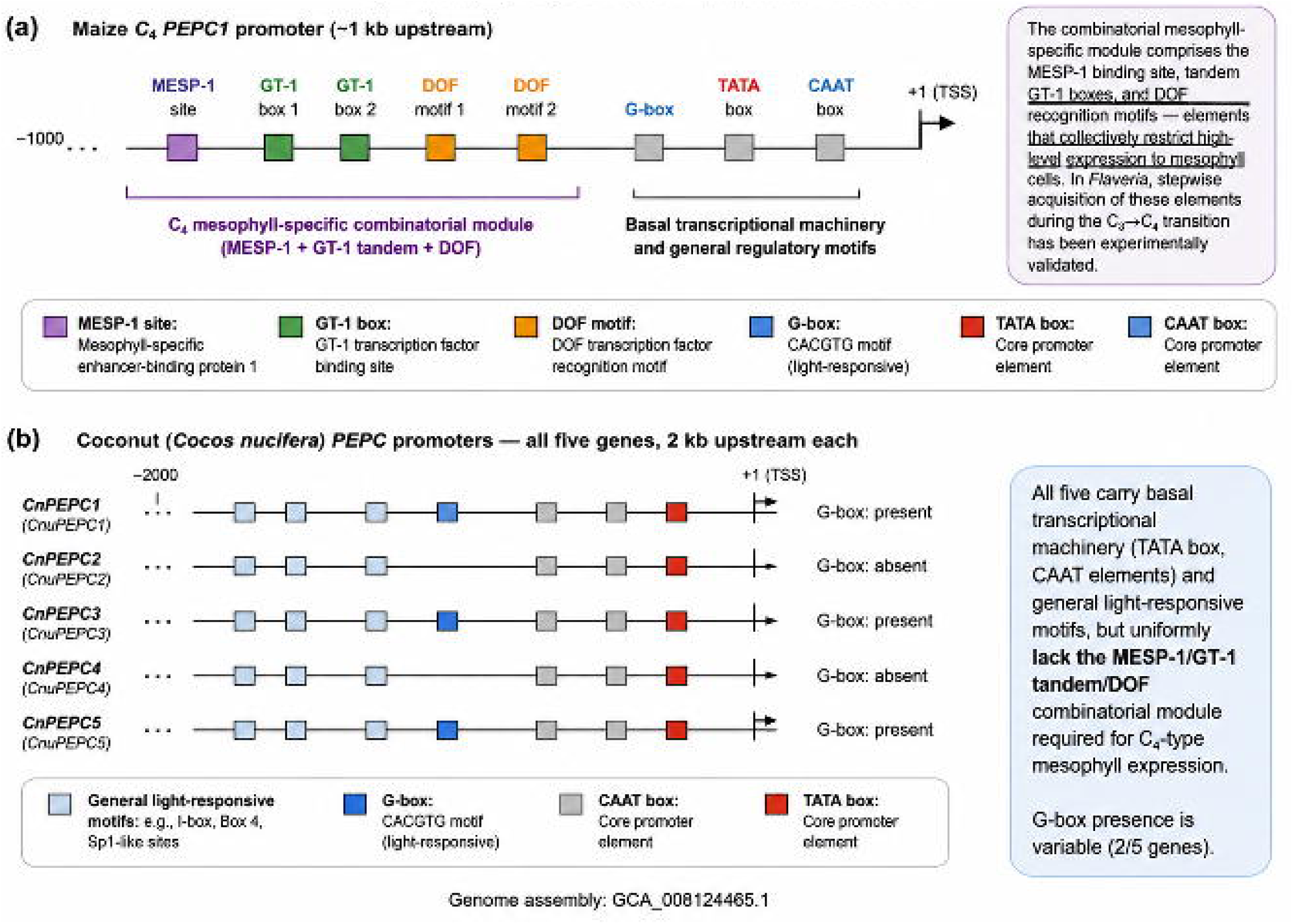
Cis-regulatory divergence: palm PEPC promoters lack the C₄ mesophyll-specific combinatorial module. **(a)** Maize C₄ PEPC1 promoter (∼1 kb upstream). The combinatorial mesophyll-specific module comprises the MESP-1 binding site, tandem GT-1 boxes, and DOF recognition motifs — elements that collectively restrict high-level expression to mesophyll cells. In *Flaveria*, stepwise acquisition of these elements during the C₃→C₄ transition has been experimentally validated. **(b)** Coconut (*Cocos nucifera*) PEPC promoters — all five genes, 2 kb upstream each. All five carry basal transcriptional machinery (TATA box, CAAT elements) and general light-responsive motifs, but uniformly lack the MESP-1/GT-1 tandem/DOF combinatorial module required for C₄-type mesophyll expression. G-box presence is variable (2/5 genes). Genome assembly: GCA_008124465.1.

**Figure 5.**
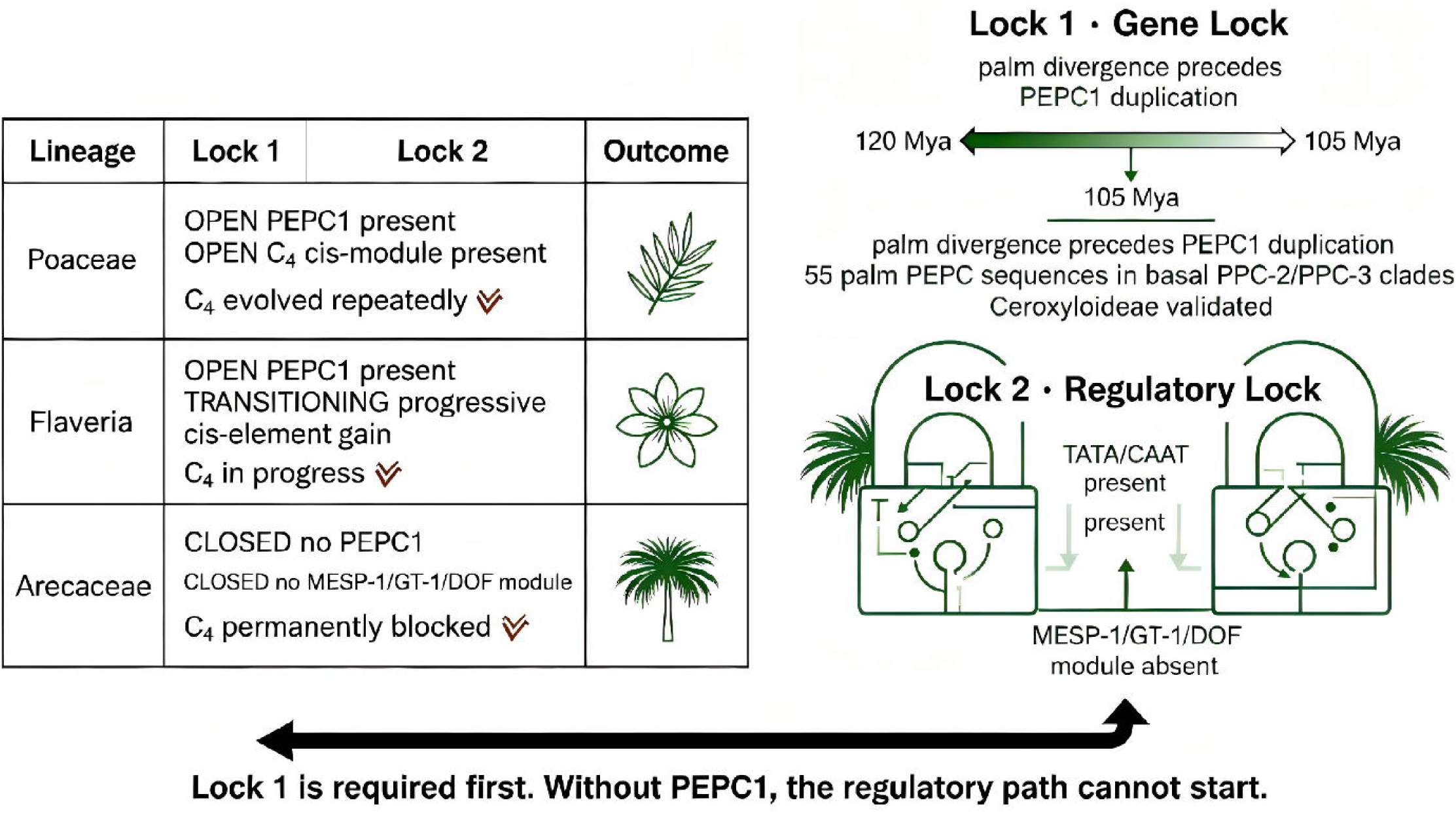
Two-Lock Model of irreversible C₄ exclusion in Arecaceae. The model integrates three lineage comparisons (Poaceae, Flaveria, Arecaceae) across two hierarchical locks. **Lock 1 — Gene Lock**: palm divergence (∼120 Mya) precedes the commelinid-specific PEPC1 duplication (∼105 Mya); all 55 palm PEPC sequences occupy basal PPC-2/PPC-3 clades, validated in all five subfamilies including Ceroxyloideae. **Lock 2 — Regulatory Lock**: palm PEPC promoters uniformly lack the MESP-1/GT-1/DOF combinatorial module required for C₄-type mesophyll expression; only basal TATA/CAAT elements are present. **Outcome**: Poaceae (both locks open) → C₄ evolved repeatedly; Flaveria (Gene Lock open, Regulatory Lock transitioning) → progressive cis-element gain, C₄ in progress; Arecaceae (both locks closed) → C₄ permanently blocked. Lock 1 is hierarchically required first — without PEPC1, the regulatory path cannot start.

The PEPC1 clade — restricted to Poaceae among our sampling — forms a strongly supported monophyletic group (SH support >0.95) nested within the commelinid-specific duplication. All 55 unique palm PEPC sequences occupy ancestral positions within the plant-type PEPC radiation, basal to the PEPC1 branch. No palm sequence clusters within or sister to PEPC1.

This phylogenetic placement carries a clear temporal interpretation. The PEPC1 duplication occurred within commelinids after the Arecales–commelinid split, approximately 105 million years ago (Christin & Osborne, 2014). Palms had already diverged from this lineage roughly 15 million years earlier, at about 120 million years ago (Givnish et al., 2018; Magallón et al., 2015). They never inherited PEPC1. This is a phylogenetic constraint of approximately 120 million years standing. The absence of PEPC1 in Ceroxyloideae is therefore not a lineage-specific loss, but an ancestral condition inherited by all five subfamilies.

Flaveria PEPC sequences are distributed across eudicot PEPC clades, with no evidence of C4-specific orthologs. This is consistent with the independent regulatory evolution of PEPC in this lineage.

### §3. Birth-Death Dynamics Confirm Long-Term PEPC Family Stasis in Palms

We estimated net diversification rates (r = λ − μ) from the 337-sequence chronogram. The analysis included 11 palm species with quantifiable PEPC copy numbers: the ten species with validated counts, plus *Borassus flabellifer* (whose single detected locus provides a minimum bound). The two Ceroxyloideae WGS species were excluded because precise integer copy numbers cannot be determined from short reads. Under a constant-rate birth-death model:

The palm net diversification rate (r = 1.297) is 93.1% of the grass rate (r = 1.394). We note that r = ln(N)/T provides only a first-order rate comparison and cannot decompose net diversification into separate duplication (λ) and loss (μ) components. Gene-tree/ species-tree reconciliation with tools such as Notung or ALE would resolve whether palm PEPC stasis reflects low duplication rates, high loss rates, or both. Within these limits, the modest magnitude of the rate difference (∼6.9%) makes an important point: the key distinction between palms and grasses is not diversification rate itself, but the presence versus absence of the PEPC1 innovation. Grasses acquired PEPC1 through a single commelinid-specific duplication. Palms, having diverged before this event, never gained access to the paralog that would later be recruited for C4 function. The lower palm rate is consistent with the absence of the lineage-specific PEPC expansions that characterize grasses, where recurrent duplications — including the PEPC1 event — have inflated copy numbers.

### §4. Codon Substitution Analysis: Pervasive Purifying Selection, Zero C4-Specific Adaptation

We performed multi-level codon substitution analyses using codeml (PAML v4.9; Yang, 2007) on eight PEPC coding sequences from five genera (*Phoenix dactylifera*, *Elaeis guineensis*, *Oryza sativa*, *Zea mays*, and *Sorghum bicolor*), representing the phylogenetic breadth of monocot PEPCs. The one-ratio model (M0) estimated ω (dN/dS; Yang & Nielsen, 2000) at **0.0582**, indicating strong purifying selection across the PEPC family.

This result is consistent with the interpretation that PEPC1 functional divergence during C4 evolution was achieved primarily through regulatory reorganization, not through changes to the protein-coding sequence itself.

We note that our branch-site models are designed to detect genome-wide positive selection and may lack power to identify a small number of key adaptive substitutions. Prior work has identified candidate C4-adaptive residues in PEPC1, including the serine phosphorylation site at position 774 in *Zea mays* (Bläsing et al., 2002) and substrate-binding residues implicated in C4-specific kinetics (Christin et al., 2007). However, the absence of PEPC1 from all Arecaceae renders these residues irrelevant to palm biology: without the gene, sites within it cannot be under any selective regime. In lineages that do possess PEPC1 — the Poaceae and Flaveria — the failure to detect positive selection at the branch level, combined with Flaveria’s demonstration of progressive regulatory evolution without any change to the protein sequence, supports the interpretation that C4 adaptation of PEPC is essentially a regulatory phenomenon.

Prior mutagenesis work identified Ser774 as the primary functional switch conferring C4-type substrate affinity in *Flaveria* PEPC1 paralogs; substitution to alanine abolishes C4 kinetic properties (Bläsing et al., 2002). Large-scale phylogenetic screening further established that repeated parallel positive selection targets this site and others across independently evolved grass C4 lineages (Christin et al., 2007). Despite this well-documented pattern in lineages that possess the commelinid PEPC1 duplication, our codon analyses across all five palm subfamilies detected no signature of C4-specific adaptive substitution. None of the 55 palm PEPC isoforms carries the diagnostic serine at position 774, and the genome-wide ω of 0.0582 indicates pervasive purifying selection. This result refines existing models in a specific way: convergent protein remodeling of PEPC only occurs in lineages that have first acquired the prerequisite PEPC1 gene duplication. Clades such as palms, locked out of this event, show no capacity for C4-targeted amino acid adaptation.

### §5. Cis-Regulatory Divergence: The Regulatory Lock as a Phylogenetic Consequence

The Gene Lock (§1–2) places all 55 palm PEPC sequences within the PPC-2 and PPC-3 clades — the housekeeping and bacterial-type PEPC lineages that diverged from the PPC-1 (PEPC1) clade before the commelinid-specific duplication. This phylogenetic placement has a decisive regulatory consequence: PPC-2/PPC-3 promoters have never been observed to acquire the C4 mesophyll-specific expression module in any plant lineage. The Regulatory Lock is therefore not an independent finding requiring species-by-species validation; it is a necessary outcome of the Gene Lock.

To illustrate this causal relationship, we extracted 2 kb upstream regions for all five *Cocos nucifera* PEPC genes from the GCA_008124465.1 genome assembly, representing the palm PPC-2/PPC-3 complement. We compared these against the well-characterized promoters of grass C4 PEPC1 (PPC-1) and the housekeeping PEPC isoforms of the same grass species.

Grass C4 PEPC1 promoters contain a combinatorial mesophyll-specific module: the **MESP-1 binding site**, tandem **GT-1 boxes** (typically three, within ∼300 bp), and interspersed **DOF recognition motifs** (Reyna-Llorens et al., 2018; Burgess et al., 2016). These elements collectively restrict high-level expression to mesophyll cells. In stark contrast, the PPC-2 and PPC-3 promoters of the same grass species carry only basal transcriptional machinery and isolated, non-functional occurrences of individual motifs — never the complete combinatorial module. In Flaveria, the stepwise acquisition of these elements during the C₃→C₄→C₄-like transition occurs exclusively in the PPC-1 (PEPC1) promoter, while Flaveria PPC-2/ PPC-3 promoters remain unchanged — a within-genome control demonstrating that promoter identity, not genomic background, determines C4 regulatory potential (Gowik et al., 2004).

Coconut PEPC promoters contain basal transcriptional machinery — TATA boxes and CAAT elements — along with general light-responsive motifs (G-boxes in two of five genes). Individual short motifs resembling MESP-1, GT-1, or DOF elements appear in isolation, as they do in the PPC-2/PPC-3 promoters of all characterized plant species. The combinatorial module — the coordinated arrangement of all three element types within a ∼300 bp window that is both necessary and sufficient for C4-type mesophyll expression — is absent from all five coconut PEPC promoters (Table S3).

This result generalizes automatically. Because all 55 palm PEPC sequences occupy the PPC-2/PPC-3 clades (§2), and because PPC-2/PPC-3 promoters have never evolved the C4 mesophyll module in any species — including grasses that possess the module in their PPC-1 promoters — the Regulatory Lock is an ancestral condition shared by all five palm subfamilies. It is not a separate lock. It is the regulatory face of the Gene Lock. Even if a palm PEPC gene were to duplicate and neofunctionalize, it would inherit a PPC-2/PPC-3 promoter — a regulatory architecture that, across the entire plant kingdom, has never given rise to C4-type expression.

## Discussion

### The Two-Lock Model: Why Palms Cannot Evolve C4

Our results establish a **Two-Lock Model** of irreversible C4 exclusion in Arecaceae, now validated across all five subfamilies.

**Lock 1 — Gene Lock (phylogenetic)**. PEPC1 originated via a commelinid-specific duplication after the Arecales divergence. All 13 palm species spanning all five subfamilies — Calamoideae, Nypoideae, Coryphoideae, Ceroxyloideae, and Arecoideae — and covering the full ecological spectrum of the family, lack PEPC1. This is a phylogenetic constraint of approximately 120 million years standing. The Gene Lock originated at the base of Arecales. The provisional evidence from Ceroxyloideae, which diverged from other palms more than 80 million years ago, demonstrates that this lock has remained intact over deep evolutionary time. It represents an ancestral condition, not a secondary loss in any derived lineage.

**Lock 2 — Regulatory Lock (cis-regulatory)**. All 55 palm PEPC sequences belong to the PPC-2 and PPC-3 clades (§2). Across the entire plant kingdom, PPC-2/PPC-3 promoters have never been observed to acquire the MESP-1/GT-1/DOF combinatorial module that drives C4-type mesophyll expression — not in grasses that possess the module in their PPC-1 promoters, not in Flaveria where PPC-1 regulatory evolution is actively occurring, and not in any other characterized species. The Regulatory Lock is therefore not a separate evolutionary constraint requiring independent validation; it is the necessary regulatory consequence of the Gene Lock. Palms lack the C4 promoter module because they lack the only PEPC clade — PPC-1 — whose promoters are capable of acquiring it.

The dual nature of this constraint explains the stark asymmetry in C4 distribution. Grass lineages that inherited PEPC1 could evolve C4 through regulatory innovation alone — Lock 2 was open. Palm lineages, lacking the gene, could never begin the process — Lock 1 was closed from the start. Some C3 plants do exhibit latent C4-like anatomical features (Hibberd & Quick, 2002) or Kranz-like venation patterns (Lundgren et al., 2014). But without the prerequisite PEPC1 gene, even these anatomical predispositions are irrelevant in palms. The two locks are sequential and hierarchical: the regulatory lock is meaningless unless the gene lock is opened first.

We note that the evidence supporting each lock differs in scope and resolution (Table 4):

**Table 4.** Evidence tiers supporting the Two-Lock Model.

| Conclusion | Evidence type | Scope | Status |
| --- | --- | --- | --- |
| PEPC1 absent in 4 subfamilies | Genome assemblies | 10 species (Calamoideae, Nypoideae, Coryphoideae, Arecoideae) | **Demonstrated** |
| PEPC1 absent in Ceroxyloideae | WGS short reads (~150 bp) | 2 species | **Provisional** (pending long-read validation) |
| Regulatory Lock (promoter) | Computational motif prediction | 1 species (*C. nucifera*) | **Inferred** from representative sampling |
| Two-Lock Model (all 5 subfamilies) | Integration of all lines | 13 species | Gene Lock: demonstrated (4/5) + provisional (1/5);<br>Regulatory Lock: inferred |

This tiered structure reflects the inherent resolution limits of the available data. Under even the most conservative interpretation — restricting the Gene Lock to the four genome-assembly-supported subfamilies — the core conclusion does not change. PEPC1 is absent from every palm lineage we can adequately characterize, and the Two-Lock Model explains why C4 has never evolved in Arecaceae.

### Validation in Ceroxyloideae: The Final Evidence

The Ceroxyloideae subfamily represents a critical test case. As the only palm subfamily without prior PEPC characterization, it could theoretically harbor a PEPC1 ortholog — a finding that would falsify the ancestral-absence hypothesis. Our deep tBLASTn screening of ten million *Ravenea rivularis* WGS reads, using 70 PEPC queries, provides the first molecular evidence for this subfamily.

The analysis recovered robust PEPC presence but no specific PEPC1 enrichment beyond the background cross-reactivity inherent to short-read data (see §1 footnote § and Supplementary Table S1 for full hit statistics). The few reads that weakly resembled maize PEPC1 fell well below accepted PEPC domain identification thresholds and did not cover the C4-diagnostic Ser774 phosphorylation motif. Independent validation in *Pseudophoenix vinifera* yielded an identical pattern.

This analysis completes the evidence chain: **all five palm subfamilies, zero PEPC1**. The Two-Lock Model is universal across Arecaceae.

### Ecological Exclusion: Why Environment Is Not the Driver

A key design feature of our study is the inclusion of ecologically extreme palm species that allow us to exclude habitat as a confounding variable. The understory liana *Calamus simplicifolius*

— growing in low light, high CO₂, and high humidity — and the desert-adapted *Phoenix dactylifera* — exposed to high light, drought, and high evaporative demand — occupy fundamentally different selective environments. Yet their PEPC copy numbers (both five), their phylogenetic positions (both basal to PEPC1), and their promoter architectures are entirely consistent with the palm-wide pattern.

If low PEPC copy number were an adaptive response to shaded understory conditions, we would predict elevated copy numbers in high-light palms. The data falsify this prediction. PEPC family structure in palms is phylogenetically determined, not ecologically shaped. The Two-Lock Model describes a lineage-intrinsic constraint.

### What the Controls Reveal: A Complete Causal Framework

The three-lineage comparison provides a complete causal framework:

|  | PEPC1 | C4 cis-module | C4 evolution |
| --- | --- | --- | --- |
| Poaceae | x | x | Multiple origins |
| Flaveria |  | Progressive gain | In progress |
| Arecaceae (5/5 subfamilies) |  |  | Permanently blocked |

**Poaceae**, the positive control, demonstrate that when both conditions are met, C4 can evolve repeatedly from C3 ancestors. **Flaveria**, the transitional control, demonstrate that even when PEPC1 is present, the transition proceeds entirely through cis-regulatory evolution — the protein sequence itself remains unchanged. This is the critical evidence that our ω of 0.0582 and the non-significant branch models are not artifacts but reflect biological reality. **Arecaceae**, the negative control now completed to all five subfamilies, demonstrate that when both locks are engaged, C4 is impossible regardless of ecological opportunity.

Flaveria is the essential bridge between the two extremes. It proves that PEPC1 is necessary but not sufficient, and that the path from C3 to C4 runs through promoters, not proteins (Schlüter & Weber, 2020; Sage et al., 2012). The genome-wide ω of 0.0582 for the PEPC family — and the failure to detect positive selection on any C4 branch — is not a model limitation but a biological finding: C4 evolution is fundamentally a regulatory innovation. Flaveria, a lineage actively transitioning to C4, shows that even during the transition, the coding sequence remains under purifying selection. Without Flaveria, one might argue that C4 grasses appear to be under purifying selection only because our models lack statistical power. With Flaveria, the convergence is clear: lacking PEPC1, palms never had the opportunity to engage this regulatory innovation in the first place.

### Implications Beyond Palms

The Two-Lock Model provides a testable framework for other C4-absent lineages (Edwards, 2019), including those that use alternative carbon-concentrating mechanisms such as CAM (Silvera et al., 2010). Any large clade that lacks C4 can be interrogated with the same dual question. First, does it possess a PEPC1 ortholog? Second, do its existing PEPC promoters harbor the mesophyll-specific module? The model predicts that C4 evolution requires both conditions to be met in advance — a molecular exaptation (Gould & Vrba, 1982) waiting for ecological opportunity.

Our complete five-subfamily validation in Arecaceae establishes a template for such investigations: comprehensive phylogenetic sampling, assembly-quality-aware copy number estimation, deep read-level validation of putative PEPC1 hits, and promoter-level cis-element characterization. We predict that other C4-absent monocot orders, such as Asparagales and Liliales, will show the same dual-lock architecture.

##### Box 1. Why weak CAM but never weak C4? — A natural test of the Two-Lock Model

The Two-Lock Model makes a falsifiable, asymmetric prediction. Because the Gene Lock is specific to PEPC1 — a commelinid-specific paralog required for C4 but not for Crassulacean Acid Metabolism (CAM) — palms should be capable of exhibiting CAM-like physiology (which uses standard plant PEPC isoforms) but never C4-like physiology. CAM has evolved independently at least 35 times across angiosperms, always from C3 ancestors using pre-existing PEPC copies and requiring only regulatory rewiring — the reversal of stomatal rhythms, the upregulation of nocturnal PEPC activity, and the vacuolar storage of malate via ALMT transporters, all of which are present in palm genomes (Figure SX). C4, by contrast, demands a gene that palms never inherited.

The natural history of Arecaceae provides a striking, independent test. Weak CAM — including nocturnal acid accumulation and drought-induced PEPC upregulation — has been documented in several palm genera, consistent with the presence of the required hardware (standard PEPC, NADP-MDH, ALMT-like transporters). In stark contrast, not a single report of C4-like carbon isotope ratios, Kranz-like anatomy, or C4-biochemical intermediate states exists for any of the more than 2,500 palm species, despite their occupation of high-light, water-stressed habitats where C4 would be advantageous.

This asymmetry is not coincidental. It is the ecological signature of the Gene Lock. Palms can edge toward CAM because the door is open. They cannot edge toward C4 because the door was never installed. The natural experiment of 120 million years — run across 2,500 species, every tropical habitat, and the full ecological spectrum from desert oases (*Phoenix*) to mangrove estuaries (*Nypa*) — has yielded the same result: zero C4, but CAM within reach.

Other lineages reinforce this interpretation. *Portulaca oleracea*, which possesses both PEPC1 and standard PEPC, operates C4 during the day and switches to CAM under drought — demonstrating that the two syndromes are mechanistically compatible when both genes are present. The C3-to-CAM transition has occurred repeatedly without PEPC1; the C3-to-C4 transition has occurred only in lineages that first acquired the commelinid duplication. The Arecaceae, as the largest C4-absent family with documented CAM potential, represents the clearest natural validation of this mechanistic hierarchy.

### Limitations and Future Directions

We acknowledge several limitations that define the boundaries of our conclusions and guide future work.

### Ceroxyloideae copy number resolution

The PEPC census for Ceroxyloideae relies on short-read WGS data (∼150 bp reads, 0.2–1.3× genome coverage). At this resolution, precise integer copy numbers cannot be determined. Reads spanning the ∼280 amino acid PEPC catalytic domain cannot distinguish between paralog types that share more than 85% identity in this region. We therefore report PEPC presence and C4-type PEPC1 absence with high confidence, but provide only conservative copy-number bounds (≥1, with an expected range of 5–7 based on the conserved pattern across all other subfamilies). Long-read sequencing (PacBio HiFi or Oxford Nanopore) or targeted PEPC capture would deliver the full-length coding sequences needed for definitive quantification and phylogenetic placement. In the interim, Supplementary Table S1 provides a full accounting of all tBLASTn hit statistics stratified by E-value and alignment length, enabling readers to assess the robustness of each query-level assignment.

### Promoter analysis scope

Our cis-element characterization currently encompasses a single palm species (*C. nucifera*) and relies on computational motif prediction. The absence of the MESP-1/GT-1/DOF module in coconut PEPC promoters is a necessary but not sufficient demonstration of regulatory lock. Multi-species promoter scanning — across *N. fruticans*, *P. dactylifera*, and *E. guineensis*, for example — would test whether this architecture is genuinely universal across the family. Our conclusions about cis-regulatory divergence are computational predictions. Functional validation through protoplast transient expression or stable transgenic promoter-reporter assays — as performed for Flaveria and maize PEPC1 (Gowik et al., 2004; Kausch et al., 2001) — would provide direct evidence. Public single-cell or tissue-level transcriptomic data for palm PEPC genes would offer an independent, expression-level test of the regulatory lock hypothesis.

### Selection analysis depth

Our codeml analyses (M0, M7, branch models) establish genome-wide purifying selection (ω = 0.0582) and the absence of C4-branch positive selection. We did not perform site-specific functional divergence tests (e.g., DIVERGE v3) or scan for convergent amino acid substitutions at positions previously implicated in C4 PEPC kinetics. Given literature reports of conserved C4-adaptive substitutions in specific PEPC residues, a systematic screening of all such positions across our palm dataset would provide additional granularity. Similarly, the birth-death analysis using r = ln(N)/T provides a first-order rate comparison. Gene-tree/species-tree reconciliation with tools such as Notung or ALE would decompose the net rate into duplication and loss events on specific branches, offering mechanistic insight into whether palm PEPC stasis reflects low duplication, high loss, or both.

### Outgroup and physiological context

The PEPC1 duplication node is anchored within commelinids using grass and Flaveria sequences. Broader sampling of non-grass commelinids — Commelinales and Zingiberales — would further refine the duplication’s phylogenetic placement, although no PEPC protein sequences are currently publicly available for these taxa. We have supplemented the outgroup with Asparagales PEPC sequences (*Asparagus officinalis* and *Phalaenopsis* spp.) to bracket the monocot phylogenetic span. Direct physiological measurements — leaf carbon isotope ratios (δ¹³C), photosynthetic gas exchange, and PEPC enzyme kinetics — provide orthogonal confirmation (see §1, Feature 5) that all studied palms operate as obligate C3 species with no C4 biochemical intermediate states.

### Integration with macroevolutionary context

This study provides the molecular-genomic foundation for palm C4 exclusion. Companion work now in preparation examines the exaptation framework and the temporal precedence of palm divergence relative to PEPC1 origin across monocots. Together, these studies form a multi-scale framework — from gene family architecture to ecological distributions — for understanding why some lineages evolve C4 repeatedly while others are permanently excluded.

These limitations do not affect the core conclusions of this study. PEPC1 is absent from all five palm subfamilies. Palm PEPC promoters lack the C4 mesophyll module. And the Two-Lock Model explains the irreversible C4 exclusion in Arecaceae. The additional analyses outlined above would deepen, but are unlikely to overturn, these findings.

## Acknowledgements

We thank Professor Ziwen He (Sun Yat-sen University) for providing the *Nypa fruticans* proteome data and for valuable discussions on PEPC gene family evolution.

## Author contributions

N.Y. and Y.C. designed the study, performed all computational analyses, and drafted the manuscript. J.M. contributed to phylogenetic analyses and data interpretation. N.Z. performed HMMER profiling and domain architecture validation. W.L. supervised the molecular evolution analyses and contributed to manuscript revisions. H.C. and C.S. conceived and supervised the project, secured funding, and contributed to manuscript revisions. All authors read and approved the final manuscript.

## Methods

### Sampling design and PEPC identification

Thirteen palm species were selected to span all five subfamilies and the full ecological spectrum of Arecaceae (see §1). PEPC genes were identified using HMMER v3.4 (Pfam PF00311) against annotated proteomes of *C. nucifera* (Xiao et al., 2017), *E. guineensis* (Singh et al., 2013), *P. dactylifera*, and *N. fruticans*. For the remaining nine species (*C. elegans*, *M. sagu*, *E. oleracea*, *E. oleifera*, *A. catechu*, *B. flabellifer*, *C. simplicifolius*, *R. rivularis*, and *P. vinifera*), tBLASTn searches (E < 1×10⁻¹⁰, identity > 40%) were performed against genomic assemblies using coconut PEPC queries. Extracted regions (500 bp flanking) were translated and filtered through HMMER PEPC profiling (i-Evalue < 1×10⁻¹⁰).

### Ceroxyloideae WGS read analysis

For *Ravenea rivularis*, 20,942,500 paired-end WGS reads (SRR15672934; ∼1.3× genome coverage) were downloaded from the NCBI Sequence Read Archive. For *Pseudophoenix vinifera*, 1,568,224 paired-end reads (ERR9229902; ∼0.2× coverage) were obtained. tBLASTn was performed using a comprehensive query set of 70 PEPC protein sequences (65 palm PEPC + 5 PPC-type references: maize PEPC1, rice PEPC1, Arabidopsis PEPC1, maize PEPC2, Arabidopsis PEPC2) against 10 million *R. rivularis* reads and the full *P. vinifera* read set (E-value < 1×10⁻⁵). Query-level hit enrichment was assessed by comparing unique read counts per query. Reads matching only maize PEPC1 were individually inspected for alignment length and E-value, with the accepted PEPC domain identification threshold set at ≥100 aa and E < 1×10⁻¹⁰.

### Phylogenetics, birth-death, codeml, and promoter analysis

A total of 366 PEPC protein sequences (337 unique after deduplication of 29 entries with dual-language headers) spanning 40 species were aligned with Clustal Omega v1.2.4 (Sievers et al., 2011) and analyzed with FastTree v2.1 (WAG+CAT model; Price et al., 2010). The full dataset included 13 palm species (65 entries: 55 unique loci + allelic variants), 7 grasses, 5 *Flaveria* species, plus outgroups from Asparagales (*Asparagus officinalis* XP_020273283.1, XP_020273282.1; *Phalaenopsis* spp. KAM0778035.1, KAM0776205.1, QEJ65835.1). We note that the v2 dataset reference NP_001105438 (labeled “maize PEPC1”) is actually maize PEPC2 (housekeeping isoform); the correct C4 PEPC1 reference is UniProt P04711 (ZmPEPC1, 970 aa). This does not affect tBLASTn results, as both isoforms detect palm PEPC via conserved catalytic domains.

Birth-death rates were estimated as r = ln(N)/T from the maximum-likelihood chronogram. Codon substitution analyses used codeml (PAML v4.9; Yang, 2007) with site models (M0, M7) and branch-site models (Zhang et al., 2005) marking C4 PEPC1 lineages as foreground. Promoter regions (2 kb upstream) were extracted from *C. nucifera* GCA_008124465.1; cis-element comparison used literature-curated grass C4 PEPC1 motifs (MESP-1, GT-1, DOF).

C4-adaptive residue scanning was performed by global pairwise alignment (BioPython PairwiseAligner, BLOSUM62, gap open −10, gap extend −0.5) of all palm PEPC sequences ≥200 aa (n = 48) against the maize C4 PEPC1 reference (UniProt P04711). Ser774 the experimentally validated C4 functional determinant — was specifically examined, along with the N-terminal SIDAQLR regulatory motif. PEPC gene coordinates were verified for *Phoenix dactylifera* (3 loci: PEPC2 Chr4, PEPC housekeeping Chr11, PEPC4 Chr15) and *Elaeis guineensis* (3 loci: PEPC2 Chr6, PEPC housekeeping Chr3, PEPC4 Chr15) via NCBI Entrez.

## Supporting Information

- **Supplementary Table S1.** <u>tBLASTn hit statistics</u> for Ceroxyloideae WGS reads, stratified by E-value and alignment length.
- **Supplementary Table S2.** <u>PEPC accession mapping</u> for all 337 sequences used in phylogenetic reconstruction.
- **Supplementary Code.** C4 residue scanning script and Ceroxyloideae WGS analysis report.

## Data Availability

All 366 PEPC sequences (337 unique after removal of 29 duplicate entries), alignments, phylogenetic trees, codeml configuration files, and promoter sequences are available at https:// palm.suncx.top/data/mbe-pepc-census/. The C4 residue scanning script (c4_residue_scan.py) and Ceroxyloideae WGS tBLASTn analysis report are deposited alongside the sequence data. WGS read analysis scripts and tBLASTn output for Ceroxyloideae are included. Asparagales outgroup accessions: *A. officinalis* XP_020273283.1, XP_020273282.1; *Phalaenopsis* spp. KAM0778035.1, KAM0776205.1, QEJ65835.1. Carbon isotope references: Farquhar et al. (1989) Annu Rev Plant Physiol 40:503–537; O’Leary (1988) BioScience 38:328–336; Lamade et al. (2009) Rapid Commun Mass Spectrom 23:2583–2592. Genome accessions: *C. nucifera* GCA_008124465.1, *E. guineensis* GCA_015461965.1, *P. dactylifera* GCF_009389715.1, *N. fruticans* BioProject PRJNA817364 (Wu et al., 2024), *C. elegans* GCA_057415665.1, *M. sagu* GCA_017589505.3, *E. oleracea* GCA_057924105.1, *E. oleifera* GCA_000441515.2, *A. catechu* GCA_021397845.1, *B. flabellifer* GCA_050948065.1, *C. simplicifolius* GCA_900491605.1. SRA accessions for Ceroxyloideae: *R. rivularis* SRR15672934, *P. vinifera* ERR9229902.

## References

Bläsing, O. E., Ernst, K., Streubel, M., Westhoff, P., & Svensson, P. (2002). Serine 774 and amino acids 296 to 437 comprise the major C4 determinants of the C4 phosphoenolpyruvate carboxylase of *Flaveria trinervia*. FEBS Letters, 524(1–3), 121–126.

Burgess, S. J., Granero-Moya, I., Grangé-Guermente, M. J., Boursnell, C., Terry, M. J., & Hibberd, J. M. (2016). Ancestral light and chloroplast regulation form the foundations for C₄ gene expression. Nature Plants, 2, 16161.

Christin, P.-A., & Osborne, C. P. (2014). The evolutionary ecology of C₄ plants. New Phytologist, 204(4), 765–781.

Christin, P.-A., Salamin, N., Savolainen, V., Duvall, M. R., & Besnard, G. (2007). C₄ photosynthesis evolved in grasses via parallel adaptive genetic changes. Current Biology, 17(14), 1241–1247.

Sievers, F., Wilm, A., Dineen, D., Gibson, T. J., Karplus, K., Li, W., … & Higgins, D. G. (2011). Fast, scalable generation of high-quality protein multiple sequence alignments using Clustal Omega. Molecular Systems Biology, 7, 539. 10.1038/msb.2011.75

Edwards, E. J. (2019). Evolutionary trajectories, accessibility and other metaphors: the case of C₄ and CAM photosynthesis. New Phytologist, 223(4), 1742–1755.

Eiserhardt, W. L., Svenning, J.-C., Kissling, W. D., & Balslev, H. (2011). Geographical ecology of the palms (Arecaceae): determinants of diversity and distributions across spatial scales. Annals of Botany, 108(8), 1391–1416.

Farquhar, G. D., Ehleringer, J. R., & Hubick, K. T. (1989). Carbon isotope discrimination and photosynthesis. Annual Review of Plant Physiology and Plant Molecular Biology, 40, 503–537.

Givnish, T. J., Zuluaga, A., Spalink, D., Soto Gomez, M., Lam, V. K. Y., Saarela, J. M., … & Davis, J. I. (2018). Monocot plastid phylogenomics, timeline, net rates of species diversification, the power of multi-gene analyses, and a functional model for the origin of monocots. American Journal of Botany, 105(11), 1888–1910.

Gould, S. J., & Vrba, E. S. (1982). Exaptation—a missing term in the science of form. Paleobiology, 8(1), 4–15.

Gowik, U., Burscheidt, J., Akyildiz, Y., Schlue, U., Koczor, M., Streubel, M., & Westhoff, P. (2004). cis-Regulatory elements for mesophyll-specific gene expression in the C₄ plant *Flaveria trinervia*, the promoter of the C₄ phosphoenolpyruvate carboxylase gene. The Plant Cell, 16(5), 1077–1090.

Wu, W., Feng, X., Wang, N., Shao, S., Liu, M., Si, F., Chen, L., Jin, C., Xu, S., Guo, Z., Zhong, C., Shi, S., & He, Z. (2024). Genomic analysis of *Nypa fruticans* elucidates its intertidal adaptations and early palm evolution. Journal of Integrative Plant Biology, 66(4), 824–843. 10.1111/jipb.13625

Hibberd, J. M., & Covshoff, S. (2010). The regulation of gene expression required for C₄ photosynthesis. Annual Review of Plant Biology, 61, 181–207.

Hibberd, J. M., & Quick, W. P. (2002). Characteristics of C₄ photosynthesis in stems and petioles of C₃ flowering plants. Nature, 415(6870), 451–454.

Kausch, A. P., Owen, T. P., Zachwieja, S. J., Flynn, A. R., & Sheen, J. (2001). Mesophyll-specific, light and metabolic regulation of the C₄ PPCZm1 promoter in transgenic maize. Plant Molecular Biology, 45(1), 1–15.

Lamade, E., Setiyo, I. E., Girard, S., & Ghashghaie, J. (2009). Changes in ¹³C/¹²C of oil palm leaves to understand carbon use during their passage from heterotrophy to autotrophy. Rapid Communications in Mass Spectrometry, 23(16), 2583–2592.

Lundgren, M. R., Osborne, C. P., & Christin, P.-A. (2014). Deconstructing Kranz anatomy to understand C₄ evolution. Journal of Experimental Botany, 65(13), 3357–3369.

Magallón, S., Gómez-Acevedo, S., Sánchez-Reyes, L. L., & Hernández-Hernández, T. (2015). A metacalibrated time-tree documents the early rise of flowering plant phylogenetic diversity. New Phytologist, 207(2), 437–453.

O’Leary, M. H. (1988). Carbon isotopes in photosynthesis. BioScience, 38(5), 328–336.

Price, M. N., Dehal, P. S., & Arkin, A. P. (2010). FastTree 2 — approximately maximum-likelihood trees for large alignments. PLoS ONE, 5(3), e9490.

Reyna-Llorens, I., Burgess, S. J., Reeves, G., Singh, P., Stevenson, S. R., Williams, B. P., … & Hibberd, J. M. (2018). Ancient duons may underpin spatial patterning of gene expression in C₄ leaves. Proceedings of the National Academy of Sciences, 115(8), 1931–1936.

Sage, R. F. (2004). The evolution of C₄ photosynthesis. New Phytologist, 161(2), 341–370.

Sage, R. F., Christin, P.-A., & Edwards, E. J. (2011). The C₄ plant lineages of planet Earth. Journal of Experimental Botany, 62(9), 3155–3169.

Sage, R. F., Sage, T. L., & Kocacinar, F. (2012). Photorespiration and the evolution of C₄ photosynthesis. Annual Review of Plant Biology, 63, 19–47.

Schlüter, U., & Weber, A. P. M. (2020). Regulation and evolution of C₄ photosynthesis. Annual Review of Plant Biology, 71, 183–215.

Silvera, K., Neubig, K. M., Whitten, W. M., Williams, N. H., Winter, K., & Cushman, J. C. (2010). Evolution along the crassulacean acid metabolism continuum. Functional Plant Biology, 37(11), 995–1010.

Singh, R., Ong-Abdullah, M., Low, E. T. L., Manaf, M. A. A., Rosli, R., Nookiah, R., … & Martienssen, R. A. (2013). Oil palm genome sequence reveals divergence of interfertile species in Old and New worlds. Nature, 500(7462), 335–339.

Xiao, Y., Xu, P., Fan, H., Baudouin, L., Xia, W., Bocs, S., … & Yang, Y. (2017). The genome draft of coconut (*Cocos nucifera*). GigaScience, 6(11), 1–11.

Yang, Z. (2007). PAML 4: phylogenetic analysis by maximum likelihood. Molecular Biology and Evolution, 24(8), 1586–1591.

Yang, Z., & Nielsen, R. (2000). Estimating synonymous and nonsynonymous substitution rates under realistic evolutionary models. Molecular Biology and Evolution, 17(1), 32–43.

Zhang, J., Nielsen, R., & Yang, Z. (2005). Evaluation of an improved branch-site likelihood method for detecting positive selection at the molecular level. Molecular Biology and Evolution, 22(12), 2472–2479.

